# Serial dependence carries the action that produced the previous report

**DOI:** 10.64898/2026.10.27.762257

**Authors:** Chunyu Qu, Corinna Naeve, Zhuanghua Shi

## Abstract

Serial dependence pulls current judgments toward the recent past, but what is carried forward is debated: the previous stimulus, the previous decision, or something about the report itself. Three magnitude-reproduction experiments separated these sources. Participants viewed dot arrays that varied independently in duration and numerosity, then reproduced the cued dimension by holding a key while a display accumulated, so that both magnitudes were produced through the same action. A response-history model partitioned the attraction into previous-stimulus and previous-response paths. The previous response, after controlling for the previous stimulus, carried the attraction across dimensions: a numerosity report biased the next duration judgment and vice versa (Experiment 1), also when each task used its own response key (Experiment 2). A task switch weakened this carryover without abolishing it. Experiment 3 randomized, trial by trial, whether a reported number required a long or a short key hold. The same previous number lengthened the next duration judgment after a long hold but not after a short one, so the carrier was the production of the report rather than its value. Conventional stimulus-based measures showed none of this cross-task history. Serial dependence can therefore carry the trace of the action through which a previous judgment was expressed, bounded but not gated by task context.

---

Judgments of quantity lean toward the judgment that came before. In the laboratory, this pull of the recent past is known as serial dependence (Cicchini et al., 2024; J. Fischer & Whitney, 2014; Manassi et al., 2023), and its source is usually sought in what was represented: the previous stimulus, or the previous decision. Yet no estimate is observed directly. It becomes visible through an action. A key is pressed, a cursor is moved, a display is adjusted. Each of these acts takes time, and each leaves a trace of its own. The experiments reported here ask whether that trace, and not only the content it delivered, shapes the judgment that follows.

Studies that separate stimulus history from response history point toward the response. A previous stimulus can repel the current percept through adaptation, whereas information retained for a later decision or report can attract the next judgment (Bansal et al., 2023; Bliss et al., 2017; Czoschke et al., 2019; Fritsche et al., 2017; Pascucci et al., 2019). When both influences enter one model, judgments are often repelled from the stimulus yet attracted toward the preceding decision or report (Fritsche et al., 2017; Gallagher & Benton, 2024; Pascucci et al., 2019; Sadil et al., 2024). Reporting also strengthens the attraction. In numerosity estimation it is stronger after a stimulus that was reported than after one that was only viewed (Morimoto & Makioka, 2024), and in duration reproduction it follows active reproduction but not passive viewing (Cheng, Chen, Glasauer, et al., 2024). Comparable effects of the previous response appear in other judgments of numerosity and duration (Fornaciai & Park, 2018; Wehrman et al., 2020; Wiener et al., 2014).

Response history also depends on the task. When two features are encoded but only one is reported, later attraction follows the reported feature (Bae & Luck, 2020), and temporal carryover is stronger after a duration judgment than after a direction judgment of the same display (Cheng, Chen, & Shi, 2024). In these designs, however, the task and the reporting action change together, and the action matters in its own right. Switching the hand that gives the response weakens the serial dependence of confidence ratings (Bocheva & Rahnev, 2025). When the same duration is reported either by continuous reproduction or by a short-or-long choice, carryover remains within each response mode and disappears across modes (Wu, Serçe, et al., 2026). A reduction after a task switch could therefore come from the task, the decision, or the motor act that delivers the report.

A second question concerns how far response history travels across magnitude dimensions. Time, space, and number have been proposed to share a generalized magnitude system (Bueti & Walsh, 2009; Walsh, 2003), although their processing dissociates in many studies of children and adults (Hamamouche & Cordes, 2019). Serial dependence offers a test of this overlap. Togoli, Fedele, et al. (2021) found attraction only within the task-relevant dimension: numerosity biased later numerosity judgments, and duration biased later duration judgments. They read the absence of transfer as evidence for dimension-specific codes. Fornaciai et al. (2026) later found serial dependence between duration and numerosity in sequences of dot arrays, and its strength tracked how strongly the two magnitudes were integrated within a stimulus. They concluded that serial dependence arises from computations shared across dimensions, which combine past and present information into one estimate of magnitude. Task-irrelevant symbolic numerals likewise attract numerosity judgments of dot arrays (Fornaciai & Park, 2022), consistent with an abstract magnitude code.

Togoli, Fedele, et al. (2021) and Fornaciai et al. (2026) both locate serial dependence in what is represented, and neither design reported both magnitudes. Observers judged one dimension per session or condition, by discrimination in the former study and by reproduction in the latter, so the other magnitude was seen but never reported. A shared code can explain how a past magnitude comes to bias a different dimension. It does not explain why the same past magnitude influences some judgments and leaves others unaffected. Decisional carryover between duration judgments occurs only when the previous and current stimuli share a sensory modality (Li et al., 2023), although duration is the judged magnitude in both. Together with the response-mode boundary described above, such limits point to retrieval. A past magnitude may shape the next judgment when the current trial retrieves the episode in which that magnitude was judged and reported.

Binding and Retrieval in Action Control [BRAC; Frings et al. (2020)] describes such a mechanism. BRAC treats each trial as an event file that binds the stimulus, the task, and the response (Hommel et al., 2001; Hommel, 2004). Features that recur on a later trial retrieve the earlier event file, including its response. Retrieval is selective. In cued visual search, for example, earlier target locations affected performance only when the task repeated, whereas bindings between location and response were largely unaffected by task preparation (Toledano et al., 2026). BRAC has mainly been tested with response times and errors. The present experiments ask whether a continuously produced magnitude also enters a bound report episode, and whether retrieving that episode biases the next magnitude judgment.

These findings yield four accounts with distinct predictions. A *dimension-specific account* holds that response history stays within the judged magnitude (Togoli, Fedele, et al., 2021): carryover within duration and within numerosity, with none between them. A *shared-magnitude account* allows transfer through a common code for “more” and “less” (Fornaciai et al., 2026; Walsh, 2003). If cross-dimensional carryover depends only on a shared magnitude, the influence of a previous report should not depend on how that value was produced. A *strict task-set account* permits response history only when the same judgment is repeated, so a task switch should abolish it. A *binding-and-retrieval account* extends BRAC to magnitude reports: the reported value and the action that produced it belong to the event file, and the next trial retrieves that file to the extent that its features recur. This account makes three predictions. Response history should cross between dimensions when both are reported through the same kind of action. A task switch should weaken it, because fewer features of the episode recur, without abolishing it. Finally, the influence of a previous report should change when the relation between the reported value and its production changes.

Testing these predictions requires a report that is the same for both magnitudes. In the studies above, the two dimensions were reported through different operations: a side choice, a slider, or a key hold. How the magnitudes are presented matters as well. Within a single trial, the direction of interference varies with the task and stimulus construction (Dormal et al., 2006; Javadi & Aichelburg, 2012; Xuan et al., 2007). Togoli, Fornaciai, et al. (2021) showed that this asymmetry depends on stimulus construction. When numerosity is represented over time as the average across a sequence of dot arrays, the interference becomes more symmetric, consistent with evidence accumulation as a mechanism shared by the two magnitudes (Lambrechts et al., 2013). We therefore used an accumulate-and-stop report. Both magnitudes are produced through one time-extended action, while the encoding array stays static, so any encoding asymmetry remains in place where it can be measured.

Three experiments used this procedure. Participants viewed static dot arrays whose duration and numerosity varied independently, then received a cue to reproduce either duration (Time) or numerosity (Dots). For both tasks, participants held a key while a dot display changed and released it when the elapsed time or the visible count matched memory. Pure blocks repeated one task, whereas mixed blocks interleaved both, which allowed us to compare carryover after task repetitions and switches. We quantified conventional serial dependence from the previous stimulus and, following Wu et al. (2026), used structural equation modeling (SEM) to estimate response history while accounting for the previous and current stimuli.

Each experiment targeted a different prediction. Experiment 1 used one response key for both tasks to test whether response history crossed between duration and numerosity and whether it persisted after task switches. Experiment 2 assigned each task its own key. This made task-specific response selection observable and tested whether carryover required repetition of the same physical key. In both experiments, dots accumulated during reproduction, so a larger reported number also required a longer hold. Experiment 3 therefore randomized, on each trial, whether the visible count rose from zero or fell from 44 dots, which reversed the relation between reported number and hold duration. This manipulation tested the prediction that separates the shared-magnitude and binding-and-retrieval accounts.

## Experiment 1

Experiment 1 used one response key for both tasks. We first measured conventional serial dependence from the preceding stimulus, then estimated response history with the previous and current stimuli controlled, and finally asked whether task switching weakened it.

## Method

### Participants

Participants were recruited from the local university participant pool. All reported normal or corrected-to-normal vision and no color-vision deficiency. Target sample size was set a priori from comparable post-cued magnitude-reproduction studies, which reported attractive serial dependence of about *d*_z_ = 0.68 to 0.75 (Cheng, Chen, Yang, et al., 2024; Wu, Serçe, et al., 2026). For the smallest effect (*d*_z_ = 0.68), 19 participants give 80% power (two-tailed α = .05). We set the target at 20 valid participants, and this rationale also guided Experiments 2 and 3. The final sample included 20 participants (22 were recruited and two were excluded, M = 24.70 years, *SD* = 2.72, range 21–31; 12 female, 8 male).

The study was approved by the Ethics Committee of the Department of Psychology, LMU Munich, on 06.04.2022. Participants were informed about the study, gave written consent before participating, and received €9 per hour. The recruitment, eligibility, consent, and compensation procedures also applied to Experiments 2 and 3.

### Apparatus and stimuli

The task was programmed in PsychoPy 2025.2.1 and presented on a VPixx monitor at 1920 × 1080 resolution. The monitor was viewed from approximately 82 cm and subtended about 40° × 23°. Participants responded on a standard keyboard. Each encoding display contained circular dots (0.5° in diameter), whose positions were drawn independently and uniformly within a 15° × 15° square centered on fixation. Positions within 1.5° of the fixation were redrawn so that the fixation cross remained clear. Dot positions were generated anew for each display. Numerosity had six levels: 12, 16, 20, 24, 28, and 32 dots. Duration had also six levels from 0.6 to 1.6 s in steps of 0.2 s. Duration and numerosity varied independently.

Encoding dots were light gray on a black background. Reproduction dots were cyan (RGB 136, 204, 238) for Time and orange for Dots. Colorimeter measurements yielded luminances of 103.3 cd/m² for the gray dots (CIE *x* = .338, *y* = .380), 110.6 cd/m² for the cyan dots (*x* = .270, *y* = .338), 70.9 cd/m² for the orange dots (*x* = .517, *y* = .423), 208.0 cd/m² for the fixation cross (*x* = .330, *y* = .356), 1.5 cd/m² for the black background (*x* = .367, *y* = .362), and 7.6 cd/m² for the surround (*x* = .398, *y* = .384).

### Design and procedure

The experiment used a 6 (duration) × 6 (numerosity) × 2 (task: Time vs. Dots reproduction) within-participant design across 12 blocks: four pure Time, four pure Dots, and four mixed. Pure blocks contained 36 trials (the full 6 × 6 crossing), and mixed blocks contained 72, presenting each combination once with a Time cue and once with a Dots cue. Each participant completed 576 main trials (144 pure Time, 144 pure Dots, and 288 mixed). Block and trial order were randomized for each participant. An 18-trial practice block preceded the main task.

Both tasks used the down-arrow key: a cyan “T” cued Time reproduction and an orange dot cued Dots reproduction. Each trial started with an 800-ms fixation cross and a 100-ms blank, followed by the dot array for the target duration. In mixed blocks, the tested dimension was unknown until a retro-cue appeared after a 500-ms blank. Participants encoded both dimensions. They began reproduction by pressing and holding the response key (Figure 1). This accumulate-and-stop report was chosen so that both magnitudes were produced through the same time-extended action; the encoding array itself was static. During reproduction, the visible dot count changed in random 1–3-dot steps per 100 ms (existing dots fixed, new ones drawn from a pre-generated pool of 48 locations), and participants released the key when the elapsed time (Time trials) or visible count (Dots trials) matched the encoded magnitude. The count accumulated from an empty array. Hold duration and visible dot count covaried but were not deterministically linked. Reproduction was capped at 5 s, with a maximum display count of 48 dots, and the inter-trial interval was sampled uniformly between 1 and 2 s.

**Figure 1.**
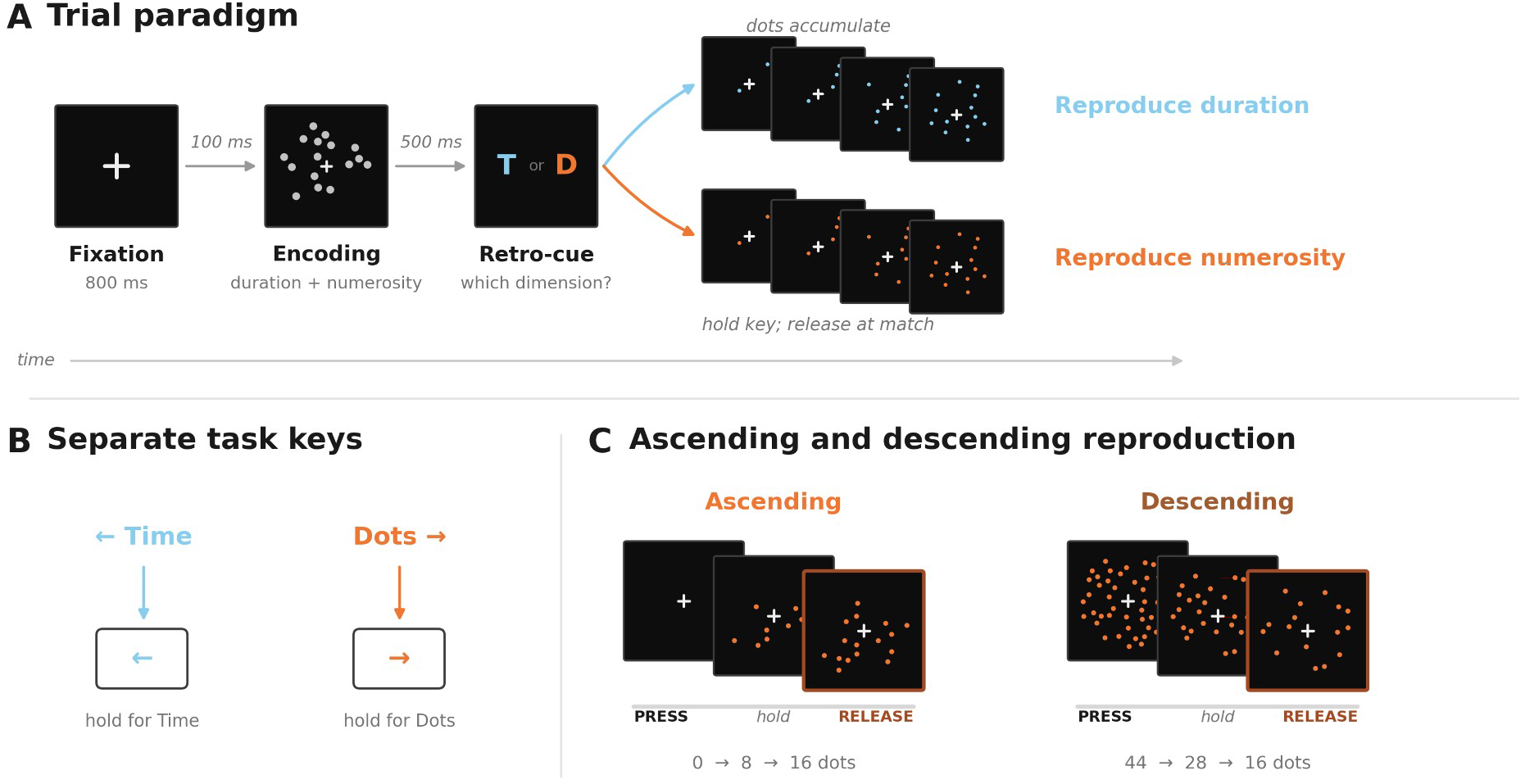
Common trial sequence and experiment-specific response manipulations. *Note.* (A) Trial paradigm. Fixation, an encoding array carrying both duration and numerosity, a retro-cue naming the dimension to reproduce, and key-hold reproduction with task-colored dots. (B) Separate task keys, used in Experiments 2 and 3; Experiment 1 used one shared key for both tasks. (C) Ascending and descending reproduction, assigned at random on every trial in Experiment 3.

Figure 1A illustrates the common trial sequence. Panels B–C preview the response manipulations introduced in Experiments 2 and 3.

## Data analysis

### Preprocessing

Means and standard deviations used to define the ±3 *SD* thresholds were calculated separately for each cue and target level within each experiment, using all practice and main trials. Practice trials were then excluded from the analysis. Participant exclusion rates were based only on outliers in the main trials, and participants with rates above 10% were removed. This threshold was set before analysis of the focal hypotheses. Lagged variables were constructed within participant and block, excluding the first trial of each block. Current bias was the reproduced value minus the target value. An analytic transition required both the current and preceding trials to be valid.

### Conventional serial dependence and response-history models

Two analyses address two questions. The first conventional serial-dependence analysis asks whether the previous stimulus attracts the current report. It shows whether our procedure reproduces the known effect and anchors the comparison with that literature. The second response-history analysis asks the question that separates the four accounts: whether the attraction is carried by the previous response and its features. A stimulus-only model cannot answer this, because the previous stimulus and the response made to it are correlated. The two analyses were therefore fitted to the same trials, and the stimulus coefficient estimated with and without the previous response shows how much of conventional serial dependence the response accounts for. The response-history model estimates carryover separately after task repetitions and task switches, so the same fit tests whether a task switch weakens it.

The conventional analysis measured the association between the previous stimulus and current reproduction bias, without reference to the previous response. For each current task, we fitted a stimulus-only regression separately to pure repetitions, mixed repetitions, and mixed switches:

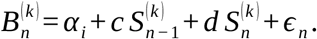

Here, 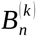 is the bias on trial *n*, calculated from the reproduced value minus target value; 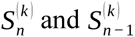 are the current and previous stimulus magnitudes, both expressed in the current judgment dimension *k*; and *α_i_* is a participant fixed effect. Positive *c* indicates attraction toward the previous stimulus after accounting for the current stimulus. On switches, the previous stimulus was duration for current Time judgments and numerosity for current Dots judgments, irrespective of the preceding cue. The coefficient therefore measures history in the current physical dimension, including when that dimension was not previously reported.

The stimulus-only regressions used the same trials as the response-history models described next. Variables were centered within participant × context and then standardized within context, so *c* is a standardized slope. Because each participant contributed many trials, standard errors were clustered by participant. With 20 participants per experiment the number of clusters is small, so we used cluster-robust standard errors with the small-sample CR2 correction and Satterthwaite degrees of freedom; all regression-based tests reported below use this combination. Each experiment yielded six tests of *c* (two tasks × three contexts), and *p* values were Holm-adjusted within this set of six. Complete estimates appear in Supplementary Section S1 (Figure S1).

The conventional coefficient cannot separate the previous stimulus from the response made to it. When the task repeats, the previous report tracks the previous stimulus, so the two sources are confounded in *c*. When the task switches, the stimulus-only model has no term for the previous response at all, although that response was produced through the same key hold and carried a value in the current dimension: a numerosity report has a hold duration, and a duration report ends at a dot count. The response-history path model therefore added the previous response and asked whether current bias is pulled toward what the participant produced on the previous trial, once the previous and current stimuli are taken into account. On repetitions, this response feature is the previous reported value. On switches, it is the feature of the previous response that corresponds to the current judgment dimension. Figure 2 shows the model for a Dots-to-Time switch, where that feature is the hold duration of the previous numerosity report, together with the Experiment 3 split of the previous response into its reported value and non-reported dimension.

**Figure 2.**
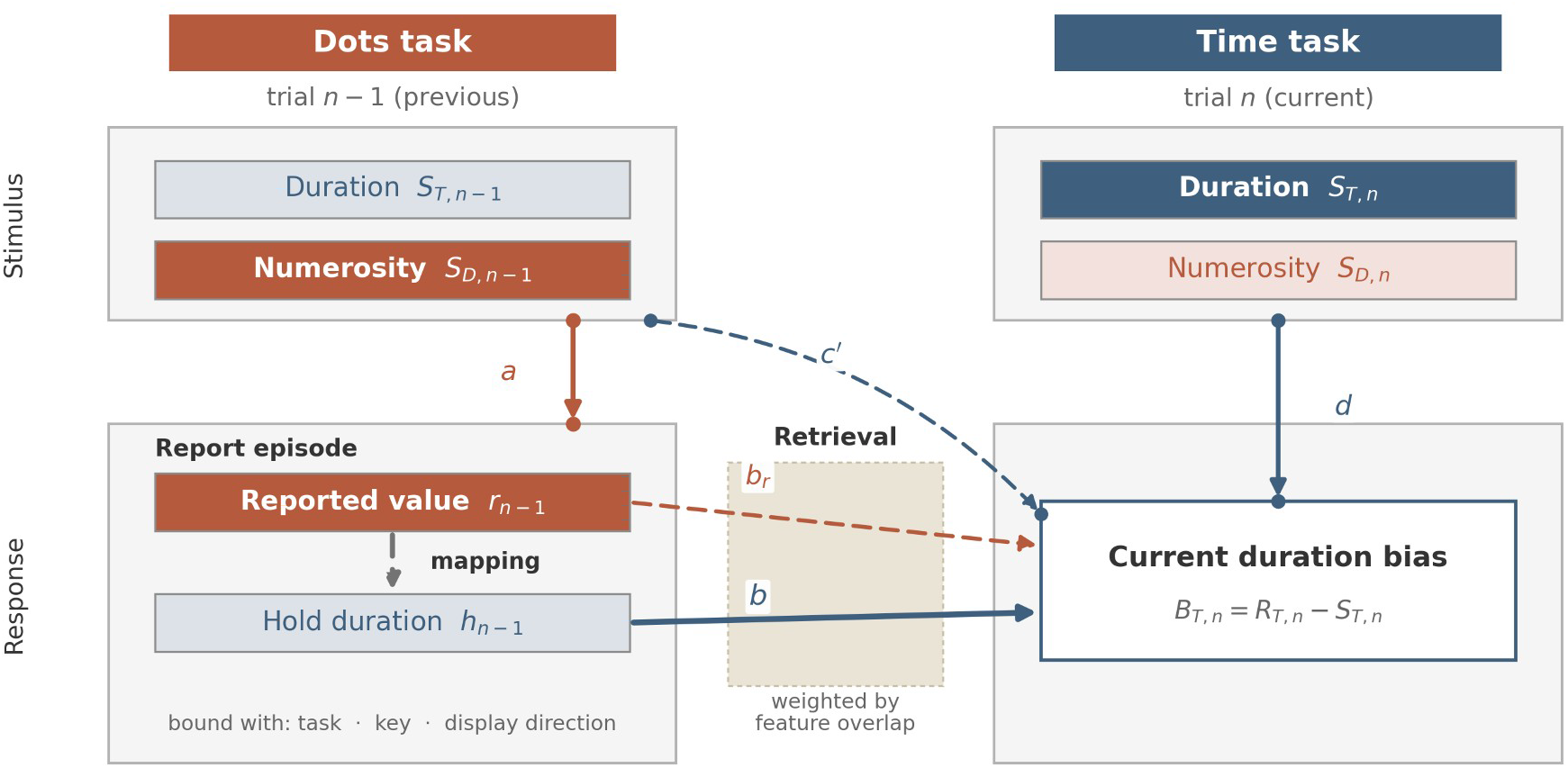
Response-history path model. *Note.* Two consecutive trials, shown for a Dots-to-Time switch. The previous report episode holds two readouts of one act: the reported value, *r_n−_*_1_, and the non-reported dimension that produced it, *h_n−_*_1_. The mapping between them was fixed in Experiments 1 and 2 and randomized in Experiment 3. Path *a* links the previous target to its report. Path *b* links the previous response feature expressed in the current dimension, here *h_n−_*_1_, to current bias; this is the response-feature carryover path of all three experiments. Experiment 3’s joint model adds *b_r_* from the reported value and controls the previous target numerosity. Paths *c*′ and *d* control the previous and current stimuli in the current dimension. In the notation of the equations below, *S_T,n_* and *S_T, n−_* _1_ are the current-dimension stimuli 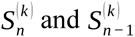 for current task *k* = Time, *S_D, n−_* _1_ is the previous-task stimulus 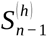, *r_n−_* _1_ is the previous report 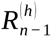 , and *h_n−_*_1_ is the response feature 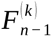. The shaded band marks retrieval of the previous episode; the transition-context and mapping analyses estimate carryover separately by the features that recur (task, block, key, display direction). Mapping interactions and participant intercepts are omitted.

One model was fitted per current task, with the previous trial sorted into three transition contexts: pure repetitions (the task repeated within a pure block), mixed repetitions (the task repeated within a mixed block), and mixed switches (the task changed within a mixed block). For previous task *h* and current task *k*, the equations were

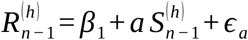

and

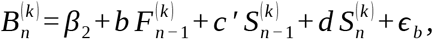

*R* denotes the previous report, *S* the stimulus, *F* the feature of the previous response expressed in the current judgment dimension, and *B* current reproduction bias. Path *a* measures stimulus-report fidelity. The key path, *b*, measures *response-feature carryover*: attraction toward the previous response feature after adjustment for the previous and current stimuli. Paths *c*′ and *d* adjust for the previous and current stimuli. Paths *a* and *b* were estimated separately in each transition context, whereas *c*′, *d*, and the covariance between the previous and current stimuli were held equal across contexts.

For a Dots-to-Time switch, *a* links previous target numerosity to reported numerosity, whereas *b* links the hold duration of that report to current Time bias. For a Time-to-Dots switch, *a* links previous target duration to reproduced duration, whereas *b* links the final dot count of that report to current Dots bias (Figure 2). Each variable was centered within participant × context and then standardized within context, so *a*, *b*, *c*′, and *d* are standardized slopes, the values plotted in Figures 3–5; each model used all complete cases. Path estimates and confidence intervals came from SEMs fitted using maximum likelihood with robust standard errors (MLR), clustered by participant. The context-specific fidelity paths and the constrained stimulus paths are reported in Supplementary Section S2 (Figure S2).

**Figure 3.**
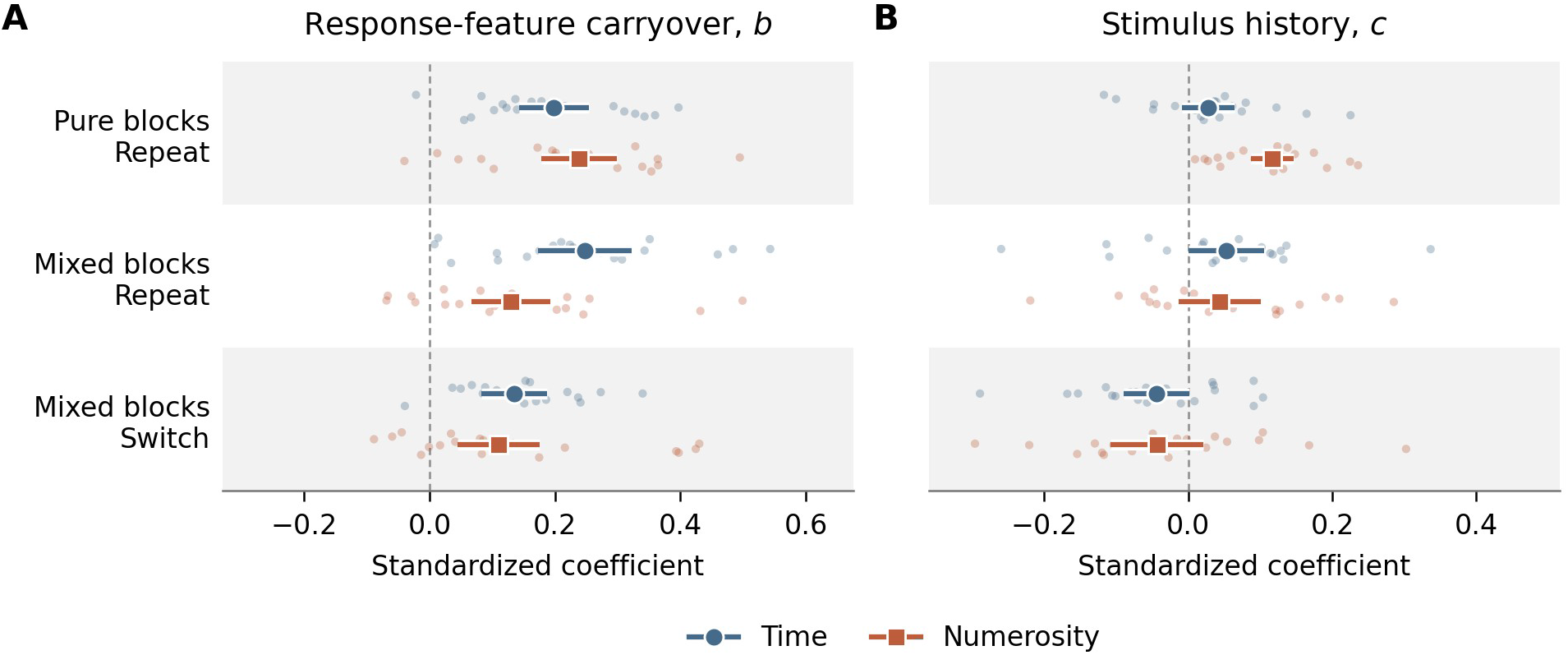
Experiment 1: Stimulus history and response-feature carryover. *Note.* Panel A shows response-feature carryover *b* and Panel B stimulus history *c*. Within each panel, rows are transition contexts, and each row shows Time (circles) above Numerosity (squares). Large points are group-model coefficients with pointwise 95% CIs (clustered MLR for *b*; CR2 for *c*). Faint dots are individual participant slopes (*n* = 20) fitted to the same trials and context scales; group points need not equal their mean. The *b* model controls both stimuli, with control slopes held equal across contexts within each fit; the *c* model controls the current stimulus. Axes are fixed by panel across Figures 3–5. Supplementary Figures S1 and S3 report the corresponding tests.

**Figure 4.**
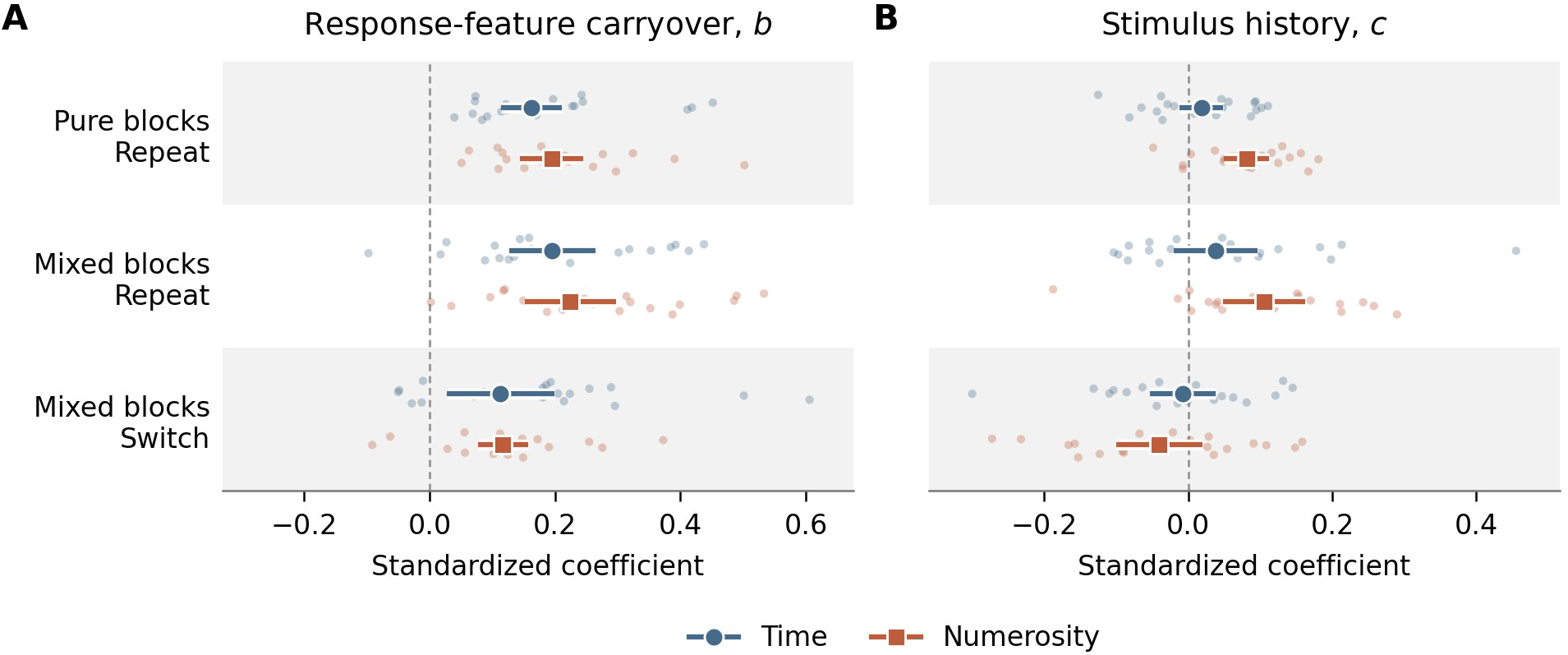
Experiment 2: Stimulus history and response-feature carryover. *Note.* Panel A shows response-feature carryover *b* and Panel B stimulus history *c*. Within each panel, rows are transition contexts, and each row shows Time (circles) above Numerosity (squares). Large points are group-model coefficients with pointwise 95% CIs (clustered MLR for *b*; CR2 for *c*). Faint dots are individual participant slopes (*n* = 20) fitted to the same trials and context scales; group points need not equal their mean. The *b* model controls both stimuli, with control slopes held equal across contexts within each fit; the *c* model controls the current stimulus. Axes are fixed by panel across Figures 3–5. Supplementary Figures S1 and S3 report the corresponding tests.

**Figure 5.**
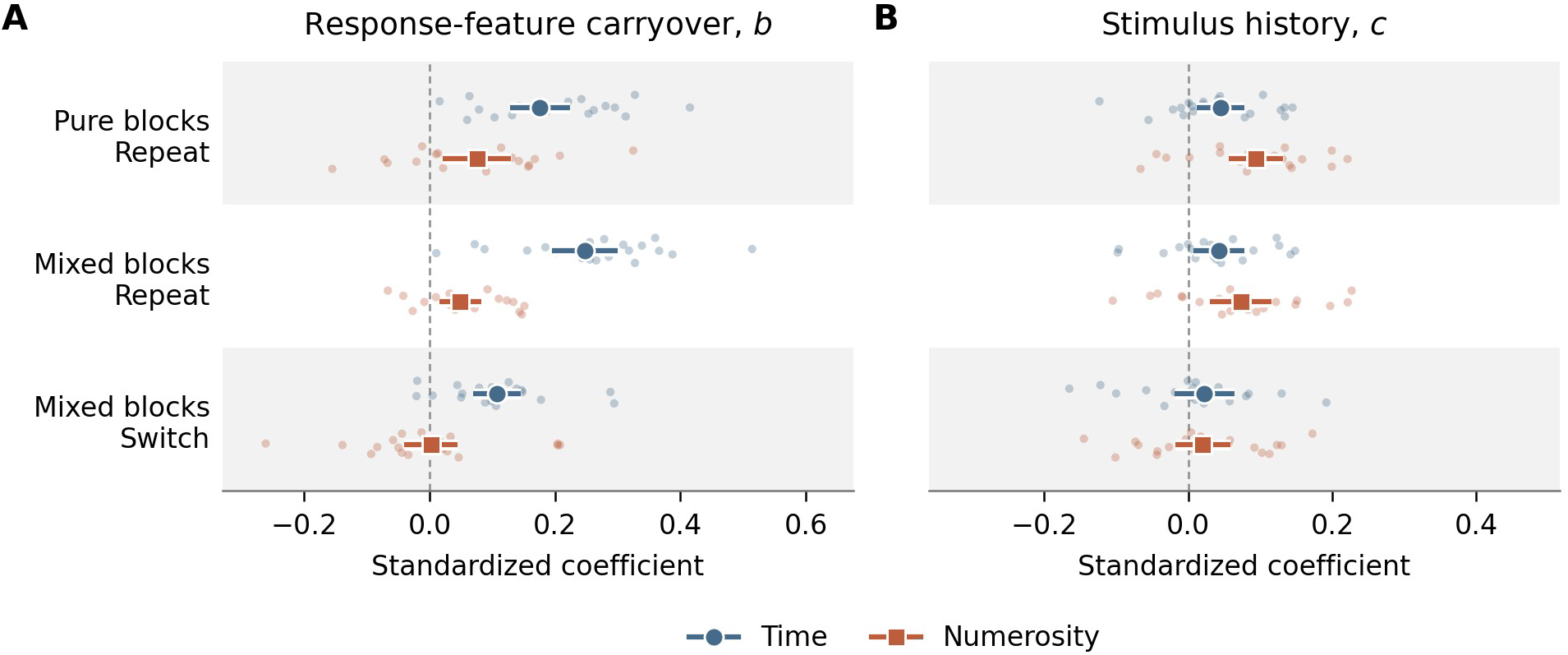
Experiment 3: Stimulus history and response-feature carryover. *Note.* Panel A shows response-feature carryover *b* and Panel B stimulus history *c*. Within each panel, rows are transition contexts, and each row shows Time (circles) above Numerosity (squares). Large points are group-model coefficients with pointwise 95% CIs (clustered MLR for *b*; CR2 for *c*). Faint dots are individual participant slopes (*n* = 20) fitted to the same trials and context scales; group points need not equal their mean. The *b* model controls both stimuli, with control slopes held equal across contexts within each fit; the *c* model controls the current stimulus. Axes are fixed by panel across Figures 3–5. Supplementary Figures S1 and S3 report the corresponding tests.

Path *c*′ is the stimulus-history association that remains once the previous response is taken into account. Comparing it with the conventional *c* therefore shows how much of conventional serial dependence is accounted for by the previous response. To compare the two on an equal footing, we also fitted the bias equation as a regression within each transition context, using the same trials, centering, and clustered inference as the stimulus-only regressions, with Holm adjustment over its own six tests per experiment. We refer to these as the matched response-adjusted regressions. Their estimates appear beside the conventional estimates in Supplementary Section S1 (Figure S1).

### Transition-context contrasts

Because *b* takes a separate value in each context, the same model tests whether carryover weakened when the task changed. The primary comparison was mixed switch versus mixed repetition, because both contexts come from mixed blocks and differ only in whether the task changed. Mixed versus pure repetition isolates block context with the task held constant. Contrasts between contexts were tested in regressions equivalent to the SEM, with participant fixed effects, participant-clustered CR2 covariance, and Satterthwaite degrees of freedom; each experiment’s six task-by-contrast tests formed one Holm family (Supplementary Section S3). All tests were two-sided. Analyses used R 4.5.2, lavaan 0.6-21, and clubSandwich 0.7.0.

The participant dots in Figures 3–5 show the spread of individual slopes around the group estimate. Because the group model yields no participant-level slopes, the bias equation was refitted for each participant on that participant’s trials by ordinary least squares, which for a single observed-variable equation equals the maximum-likelihood fit. These fits used the same complete cases and context standardization as the group model, with *b* free by context and the stimulus-control slopes shared across contexts; the stimulus-only coefficient *c* was fitted within each context, controlling the current stimulus. The dots are descriptive: the group points and intervals come from the group models and are not adjusted by them.

### Transparency and openness

The following statements apply to all three experiments.

### Data and code availability

Deidentified data and analysis code will be made publicly available upon publication. The experiments and analyses were not preregistered.

### Generative artificial intelligence

Generative AI models, including ChatGPT and Claude were used for language editing and code generation. The authors reviewed and edited all generated content, checked generated code against the source data and saved results, and take responsibility for the manuscript and analyses.

## Results

### Conventional serial dependence

Conventional stimulus-history attraction was clearest for Dots judgments in pure blocks (Figure 3B), *c* = 0.117, 95% CI [0.088, 0.146], *t*(19.0) = 8.35, *p* < .001, Holm-adjusted *p* < .001. The corresponding Time estimate was positive but uncertain, *c* = 0.028, 95% CI [−0.009, 0.065], *t*(19.0) = 1.57, *p* = .134, Holm-adjusted *p* = .387. Neither mixed-repetition estimate nor either mixed-switch estimate survived correction; the switch intervals both included zero (Supplementary Section S1, Figure S1).

### Response history within and across tasks

Current reports were attracted toward the preceding response in every transition context (Figure 3A). Within a task, carryover was positive for Time after pure repetitions, *b* = 0.198, 95% CI [0.143, 0.254], and mixed repetitions, *b* = 0.248, [0.173, 0.323], and for Dots after pure repetitions, *b* = 0.239, [0.179, 0.299], and mixed repetitions, *b* = 0.130, [0.067, 0.192], all *p* < .001.

Carryover also crossed the task boundary. On Dots-to-Time switches, current Time bias increased with the duration of the preceding numerosity response, *b* = 0.135, 95% CI [0.083, 0.187], *p* < .001. On Time-to-Dots switches, current Dots bias increased with the final dot count of the preceding duration response, *b* = 0.110, [0.044, 0.175], *p* = .001. Both associations held after adjustment for the previous and current stimuli. Thus, response features carried over even when the preceding judgment concerned a different magnitude dimension.

Switching nevertheless weakened Time carryover relative to repeating the task within mixed blocks, Δ*b* = −0.113, 95% CI [−0.188, −0.037], *t*(16.6) = −3.14, *p* = .006, Holm-adjusted *p* = .027. The corresponding Dots contrast was uncertain, Δ*b* = −0.020, [−0.094, 0.055], *t*(16.9) = −0.56, *p* = .586. Dots carryover instead decreased between pure and mixed repetitions, Δ*b* = −0.109, [−0.181, −0.037], *t*(17.1) = −3.18, *p* = .005, Holm-adjusted *p* = .027. For Dots, the clearest reduction therefore concerned block context. Holm adjustment covered the six task-by-context contrasts within each experiment. The only other contrast to survive it was the Dots difference between mixed switches and pure repetitions, Δ*b* = −0.129, Holm-adjusted *p* = .013, which combines the two changes; the remaining contrasts were uncertain (Supplementary Section S3, Figure S3).

### Stimulus history after response adjustment

In the matched response-adjusted regressions, the response-adjusted Time stimulus coefficient was negative in pure blocks, *c*′ = −0.098, 95% CI [−0.146, −0.051], *t*(17.2) = −4.34, *p* < .001, Holm-adjusted *p* = .003. The other five conditional stimulus tests did not survive correction (Supplementary Section S1, Figure S1).

## Discussion

Experiment 1 established response-feature carryover within and between the two magnitude tasks, with reduced Time carryover after a switch. However, the shared key left task differentiation during reproduction difficult to assess. Experiment 2 assigned each task a different key, requiring task-specific response selection and allowing us to check whether participants followed the task cues.

## Experiment 2

Experiment 2 tested whether participants distinguished the two cued tasks and whether response-feature carryover persisted when switching tasks also required changing keys. We evaluated task-key use before examining serial dependence.

## Method

### Participants

The sample comprised 20 participants (20 were recruited and none were excluded; M = 25.60 years, *SD* = 5.00, range 21–38; 14 female, 6 male). The target sample size and recruitment procedures followed Experiment 1.

### Design and procedure

The apparatus, stimuli, and procedure were the same as in Experiment 1, with two changes. First, the six target durations ranged from 0.8 to 1.8 s in steps of 0.2 s. Second, Time and Dots reproduction used separate, counterbalanced keys, with spatial cues indicating the current task-key mapping (Figure 1B). A wrong-key attempt produced a 500-ms red warning and a 200-ms blank before the cue reappeared. This required participants to select the key associated with the cued task. The block structure and trial counts followed Experiment 1.

### Data analysis

We used the preprocessing, response-history models, and transition-context comparisons described in Experiment 1. A trial entered the carryover analysis after a valid report, whereas cue-to-press analyses excluded trials with a wrong-key attempt.

## Results

### Task-key compliance

Participants generally selected the appropriate key. Wrong-key attempts occurred on 69 of 11,520 scheduled main trials (0.60%). Correct-key cue-to-press latencies were longer after switches than repetitions for Time, 40.47 ms, 95% CI [12.58, 68.36], *t*(19) = 3.04, *p* = .007, *d*_z_ = 0.68, and Dots, 47.25 ms, [27.52, 66.98], *t*(19) = 5.01, *p* < .001, *d*_z_ = 1.12. These checks provided behavioral evidence that participants distinguished the task cues and selected the corresponding response keys. Wrong-key rates were similar for Time (0.66%) and Dots (0.54%) and across the two key counterbalances (0.57% and 0.63%), and higher on switches (1.80%) than on repetitions (0.43%) within mixed blocks (Supplementary Section S4).

### Conventional serial dependence

With separate response keys, conventional attraction again appeared for Dots repetitions (Figure 4B). It was positive in pure blocks, *c* = 0.081, 95% CI [0.050, 0.112], *t*(19.0) = 5.45, *p* < .001, Holm-adjusted *p* < .001, and on repetitions within mixed blocks, *c* = 0.105, 95% CI [0.048, 0.162], *t*(18.8) = 3.85, *p* = .001, Holm-adjusted *p* = .006. Time associations and both mixed-switch associations were uncertain after correction (Supplementary Section S1, Figure S1).

### Response history within and across tasks

Response-feature carryover persisted with separate keys (Figure 4A). Within a task, carryover was positive for Time after pure repetitions, *b* = 0.162, 95% CI [0.114, 0.211], and mixed repetitions, *b* = 0.196, [0.126, 0.265], and for Dots after pure repetitions, *b* = 0.195, [0.145, 0.245], and mixed repetitions, *b* = 0.225, [0.152, 0.297], all *p* < .001.

The two switch paths were also positive: Dots to Time, *b* = 0.113, 95% CI [0.028, 0.199], *p* = .009, and Time to Dots, *b* = 0.117, [0.077, 0.157], *p* < .001. Carryover across tasks therefore survived when the tasks used different physical keys and participants showed measurable costs of switching between them.

Both switch-minus-mixed-repeat estimates were negative, but neither survived Holm correction. The Time contrast was uncertain, Δ*b* = −0.083, 95% CI [−0.200, 0.033], *t*(15.8) = −1.52, *p* = .149, Holm-adjusted *p* = .596. The Dots contrast was nominally significant, Δ*b* = −0.107, [−0.200, −0.015], *t*(16.6) = −2.45, *p* = .026, but its Holm-adjusted *p* = .151. Experiment 2 therefore replicated carryover across tasks. Its evidence for switch attenuation did not survive correction (Supplementary Section S3, Figure S3).

### Stimulus history after response adjustment

After response adjustment, the Time stimulus coefficient was again negative in pure blocks, *c*′ = −0.086, 95% CI [−0.132, −0.040], *t*(17.8) = −3.91, *p* = .001, Holm-adjusted *p* = .006. The remaining conditional stimulus tests did not survive correction (Supplementary Section S1, Figure S1).

## Discussion

Separate keys made task-specific response selection observable, and carryover remained positive in both directions. A further ambiguity remained: as dots accumulated during reproduction, longer holds produced larger dot counts. Experiment 3 reversed this relation on randomly selected trials to distinguish carryover tied to the reported number from carryover tied to how that number was produced.

### Experiment 3

Experiment 3 tested whether carryover from a numerosity report to the next duration judgment depended on the mapping used to produce that report. We retained separate task keys and reversed the relation between reported number and hold duration on randomly selected trials. This manipulation tested whether carryover followed the reported number alone or also depended on response production.

## Method

### Participants

The final sample comprised 20 participants (21 were recruited and one was excluded, 13 females, seven males). Their mean age was 26.80 years (*SD* = 6.64, range 20–47). The target sample size and recruitment procedures followed Experiment 1.

### Design and procedure

The apparatus, encoding stimuli, target durations, block structure, and separate task keys followed Experiment 2. We randomized the value-to-hold mapping on every trial (Figure 1C). Numerosity reports counted upward from zero (ascending mapping) or downward from 44 dots (descending mapping). A larger reported number therefore required a longer hold under the ascending mapping but a shorter hold under the descending mapping. The two start values were symmetric about the design center of 22 dots; descending arrays stopped at one dot, so the display never emptied. Mapping was assigned equally often within each cued-task × target-level combination and occurred equally often in both tasks. On Time trials, the display changed in the same two directions, but elapsed hold duration defined the report.

### Data analysis

We added two focal Dots-to-Time models to the analyses described in Experiment 1. In Experiments 1 and 2, dot accumulation coupled reported numerosity with hold duration, making their contributions difficult to distinguish. The randomized mapping gives them different signatures. If the previous reported number is what carries forward, its relation to current Time bias should keep the same direction under both mappings. If the duration of the response that produced it is what carries forward, that relation should reverse when the mapping reverses, because the same number is produced by a long hold under one mapping and a short hold under the other. The diagnostic quantity is therefore the interaction between the previous report and the previous mapping. Within either mapping, reported numerosity and hold duration remained highly collinear, so identification came primarily from randomizing their relation across mappings.

The mapping-interaction model tested this prediction. It included previous reported numerosity × previous mapping × current mapping and all lower-order terms. The joint model asked a secondary question: whether Time bias was positively associated with hold duration beyond the reported number. It entered reported numerosity and hold duration simultaneously, allowed both coefficients to vary with current mapping, and compared them (Figure 2, path *b_r_*). Both models controlled previous target numerosity and current target duration. Participants differed in how the starting count shifted their reports, and with participant intercepts alone these stable differences would enter the mapping-specific slopes. Both models therefore included a separate intercept for each participant × previous mapping × current mapping cell. In the joint model, numerosity was scaled by 22 dots, and duration and current Time bias by 1.3 s. Previous and current mappings were effect coded as ascending = +1 and descending = −1. Alternative coordinates, model restrictions, and the reverse Time-to-Dots transition appear in Supplementary Section S5.

## Results

### Conventional serial dependence

Conventional attraction was present in pure blocks for both Time (Figure 5B), *c* = 0.045, 95% CI [0.012, 0.077], *t*(19.0) = 2.83, *p* = .011, Holm-adjusted *p* = .043, and Dots, *c* = 0.094, 95% CI [0.056, 0.131], *t*(19.0) = 5.18, *p* < .001, Holm-adjusted *p* < .001. Dots attraction also survived correction on mixed repetitions, Holm-adjusted *p* = .010. The corresponding Time association was nominally positive, *p* = .022, but did not survive correction, Holm-adjusted *p* = .067. Both stimulus-only switch intervals included zero (Supplementary Section S1, Figure S1).

### Response history within and across tasks

Carryover was again positive within each task (Figure 5A): for Time after pure repetitions, *b* = 0.176, 95% CI [0.128, 0.224], and mixed repetitions, *b* = 0.248, [0.196, 0.300], both *p* < .001, and for Dots after pure repetitions, *b* = 0.076, [0.021, 0.130], *p* = .006, and mixed repetitions, *b* = 0.049, [0.016, 0.082], *p* = .004. On Dots-to-Time switches, carryover remained positive, *b* = 0.107, [0.069, 0.146], *p* < .001, whereas the mapping-averaged Time-to-Dots path was near zero, *b* = 0.002, [−0.040, 0.044], *p* = .918. Dots-to-Time carryover therefore remained positive under the randomized mapping, allowing us to examine how it varied with response production.

Switching again reduced Time carryover relative to mixed repetitions, Δ*b* = −0.140, 95% CI [−0.204, −0.076], *t*(17.6) = −4.61, *p* < .001, Holm-adjusted *p* = .001. The corresponding Dots contrast was uncertain, Δ*b* = −0.047, [−0.097, 0.004], *t*(17.4) = −1.94, *p* = .068, Holm-adjusted *p* = .137 (Supplementary Section S3, Figure S3).

### Stimulus history after response adjustment

The matched response-adjusted regressions showed negative Time stimulus associations on both pure and mixed repetitions. Dots retained a positive stimulus association in pure blocks, *c*′ = 0.052, 95% CI [0.013, 0.092], *t*(18.0) = 2.81, *p* = .012, Holm-adjusted *p* = .047. The remaining Dots and switch associations were uncertain after correction (Supplementary Section S1, Figure S1).

### Carryover follows the response mapping

The randomized mapping separates the two accounts. If the reported number itself carried over, its association with current Time bias should be positive under both mappings. If the response that produced it carried over, the association should reverse, because the same number was produced by a long hold under the ascending mapping and a short hold under the descending mapping. The manipulation worked as intended: reported numerosity and hold duration correlated *r* = .922 under the ascending mapping and *r* = −.909 under the descending mapping, compared with *r* = .165 when pooled (Figure 6A).

**Figure 6.**
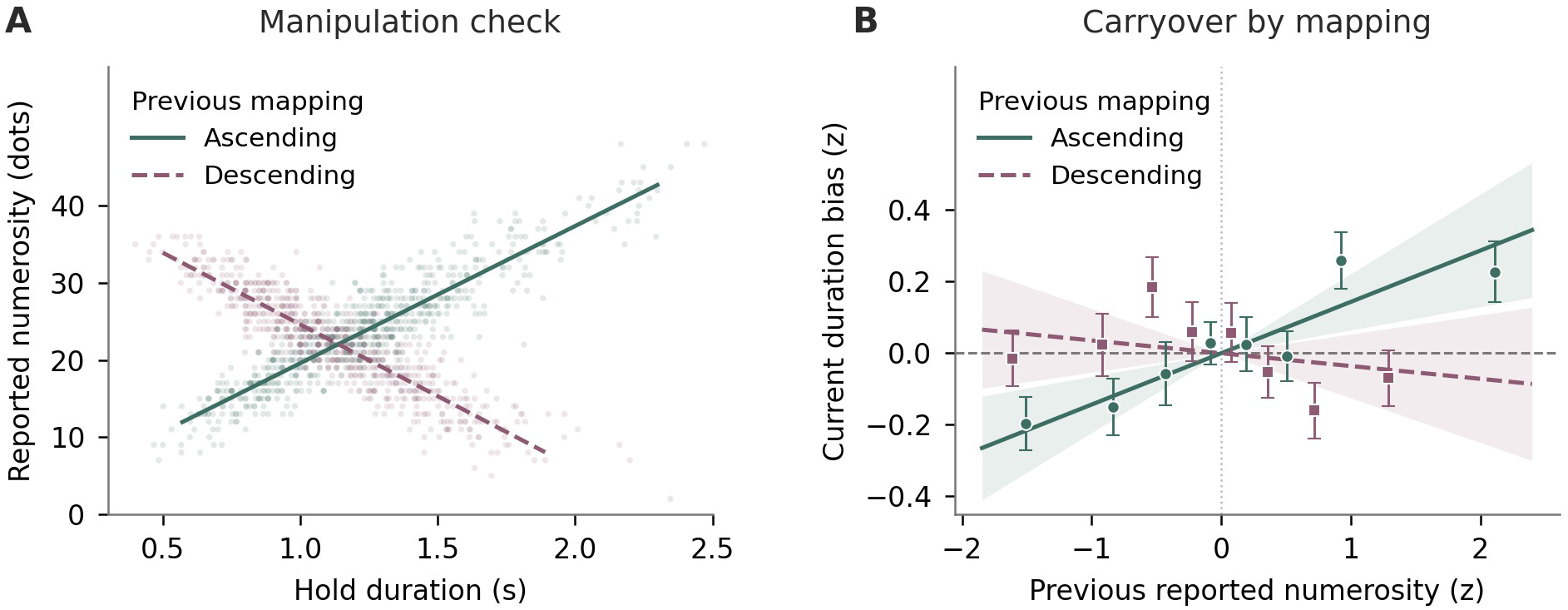
Experiment 3: Dots-to-Time carryover depends on the randomized response mapping. *Note.* (A) Manipulation check on the 1,409 Dots-to-Time switch trials. Reported numerosity and the hold duration that produced it covaried positively under the ascending mapping and negatively under the descending mapping. Lines summarize these associations descriptively. (B) Current duration bias as a function of previous reported numerosity, split by previous mapping and averaged over current mapping. Variables are participant-centered z scores. Bias is adjusted for the model’s control terms and its participant × mapping-cell intercepts, so both fitted lines pass through zero. Points show trial-level bin means and descriptive SEMs; fitted lines and bands show model estimates and participant-clustered CR2 95% CIs. Solid lines and circles denote the ascending mapping; dashed lines and squares denote the descending mapping. Joint-model coefficients are reported separately in Supplementary Section S5.

Carryover reversed with the mapping (Figure 6B). After ascending reports, current Time bias increased with the previous reported number, *b* = 0.143; after descending reports the slope was slightly negative, *b* = −0.036. The previous reported numerosity × previous mapping interaction was reliable, *b* = 0.090, 95% CI [0.051, 0.128], *p* < .001, and did not depend on the current mapping (three-way interaction *b* = −0.009, *p* = .742). Averaged over the two mappings the numerosity coefficient was uncertain, *b* = 0.054, 95% CI [−0.020, 0.128], *p* = .146, not because carryover was absent but because the two mappings pulled in opposite directions. The interaction remained reliable across alternative coordinates (Supplementary Section S5). Thus, what carried over from a numerosity report depended on how that number had been produced.

### Joint contributions of reported numerosity and hold duration

The interaction shows that carryover depended on the mapping, but not which readout of the previous response carried it. The joint model asked this directly by entering the previous reported number and the hold duration that produced it in the same model, so that each coefficient reflects one readout with the other held constant. With reported numerosity held constant, current Time bias still increased with hold duration, *b_h_* = 0.095, 95% CI [0.060, 0.131], *p* < .001. With hold duration held constant, the numerosity coefficient was uncertain, *b_r_* = 0.043, [−0.020, 0.106], *p* = .170. The difference between the two coefficients was also uncertain, Δ*b* = *b_h_* − *b_r_* = 0.052, [−0.023, 0.127], *p* = .158 (Supplementary Section S5).

Hold duration is therefore the readout with reliable independent carryover. The data do not rank the two readouts, because within each mapping they remained highly correlated. The same pattern held across alternative coordinates; a stricter fixed-effect specification was less precise (Supplementary Section S5).

### Reverse transition

The reverse Time-to-Dots transition showed no reliable carryover from the previous endpoint count. Its mapping-averaged slope was near zero, *b* = −0.010, 95% CI [−0.062, 0.043], *p* = .702, and the numerosity × previous mapping × current mapping interaction was uncertain, *b* = 0.005, [−0.050, 0.059], *p* = .850. None of the four mapping-specific slopes was reliable, whereas the previous reproduced duration, averaged over mappings, did carry over to Dots bias, *b* = 0.065, 95% CI [0.014, 0.115], *p* = .015 (Supplementary Section S5). Because reported duration and hold duration coincide on Time trials, this reverse transition could not distinguish their contributions in the same way.

## Discussion

Reversing the reproduction mapping changed the influence of a numerosity report on the next duration judgment: a larger reported number lengthened the following duration report after ascending reports, but not after descending ones. This result indicates that carryover depended on how the report was produced, beyond its numerical value alone.

## Combined analysis of task-transition attenuation

The experiment-specific comparisons showed reliable Time reproduction attenuation in Experiments 1 and 3, with less consistent evidence for Dots. A separate combined analysis summarized the evidence across all three experiments.

## Method

We fitted the response-history model described in Experiment 1 to the combined sample. Whether attenuation differed between Time and Dots was tested by fitting both tasks in a single model and contrasting the mixed-switch minus mixed-repetition effect between tasks. The six-contrast Holm sensitivity was applied to the combined context comparisons, as in the experiment-specific analyses.

The pure blocks also provide the fixed-task design of the predecessor time–number studies, with static encoding arrays. For each experiment and current task, we regressed reproduction bias in pure blocks on the current relevant and irrelevant targets (within-trial integration), and then added the previous relevant and irrelevant targets (stimulus history). These models used the complete cases of the lag-1 analyses. Variables were participant-centered and standardized within experiment and task; models included participant and block intercepts and used participant-clustered CR2 inference. Holm adjustment covered the two current tasks within each term.

## Results

Across experiments, carryover was weaker after switches than repetitions within mixed blocks. For Time judgments, the difference was Δ*b* = −0.110, 95% CI [−0.158, −0.062], *p* < .001 (mixed repetitions, *b* = 0.229, [0.190, 0.269]; mixed switches, *b* = 0.120, [0.083, 0.157]). Dots judgments showed the same direction, Δ*b* = −0.053, 95% CI [−0.094, −0.011], *p* = .015 (mixed repetitions, *b* = 0.128, [0.092, 0.165]; mixed switches, *b* = 0.076, [0.044, 0.107]). Carryover remained reliably positive on switches for both tasks (both *p*s < .001). The direct task comparison was uncertain, Δ = −0.057, 95% CI [−0.119, 0.005], *p* = .069; the interval still admitted a substantial difference in attenuation.

Block context also affected Time carryover. With the task repeated, carryover was stronger in mixed than pure blocks (pure repetitions, *b* = 0.178, 95% CI [0.147, 0.208]; Δ*b* = 0.052, [0.011, 0.093], *p* = .013). The corresponding Dots contrast was uncertain (pure repetitions, *b* = 0.165, [0.131, 0.200]; Δ*b* = −0.037, [−0.081, 0.007], *p* = .096). Mixed switches also showed weaker carryover than pure repetitions for both tasks (Time Δ*b* = −0.058, 95% CI [−0.095, −0.020], *p* = .003; Dots Δ*b* = −0.090, [−0.128, −0.051], *p* < .001), but these contrasts combine a task change with a change in block context (Supplementary Section S3, Figure S3). All five nominally reliable combined context contrasts survived the six-contrast Holm sensitivity. Figure S4 shows the trial-level trends underlying these context coefficients and those in Figures 3–5.

### Pure blocks: within-trial integration and cross-dimensional stimulus history

Within a trial, numerosity biased duration reproduction in every experiment, pooled *b* = 0.113, 95% CI [0.087, 0.140], *p* < .001 (Experiments 1–3: *b* = 0.132, 0.147, and 0.062; all Holm-adjusted *p*s < .002). Duration biased numerosity reproduction in Experiment 2 (*b* = 0.051, Holm-adjusted *p* = .022), but this effect was not evident in Experiments 1 and 3 (*b* = −0.004 and −0.032) or in the pooled estimate (*b* = 0.005, 95% CI [−0.019, 0.029]). Across trials, current bias was unrelated to the previous irrelevant magnitude once the current magnitudes and the previous relevant magnitude were controlled, pooled *b* = 0.012, 95% CI [−0.009, 0.033] for Time and *b* = 0.002, 95% CI [−0.019, 0.022] for Dots. Both intervals exclude effects larger than about 0.03, one third of the within-dimension stimulus history for Dots in the same model, *b* = 0.097, 95% CI [0.079, 0.115]. The same pattern held in each experiment: the previous irrelevant magnitude was uncertain in all six experiment-by-task cells (Figure 7).

**Figure 7.**
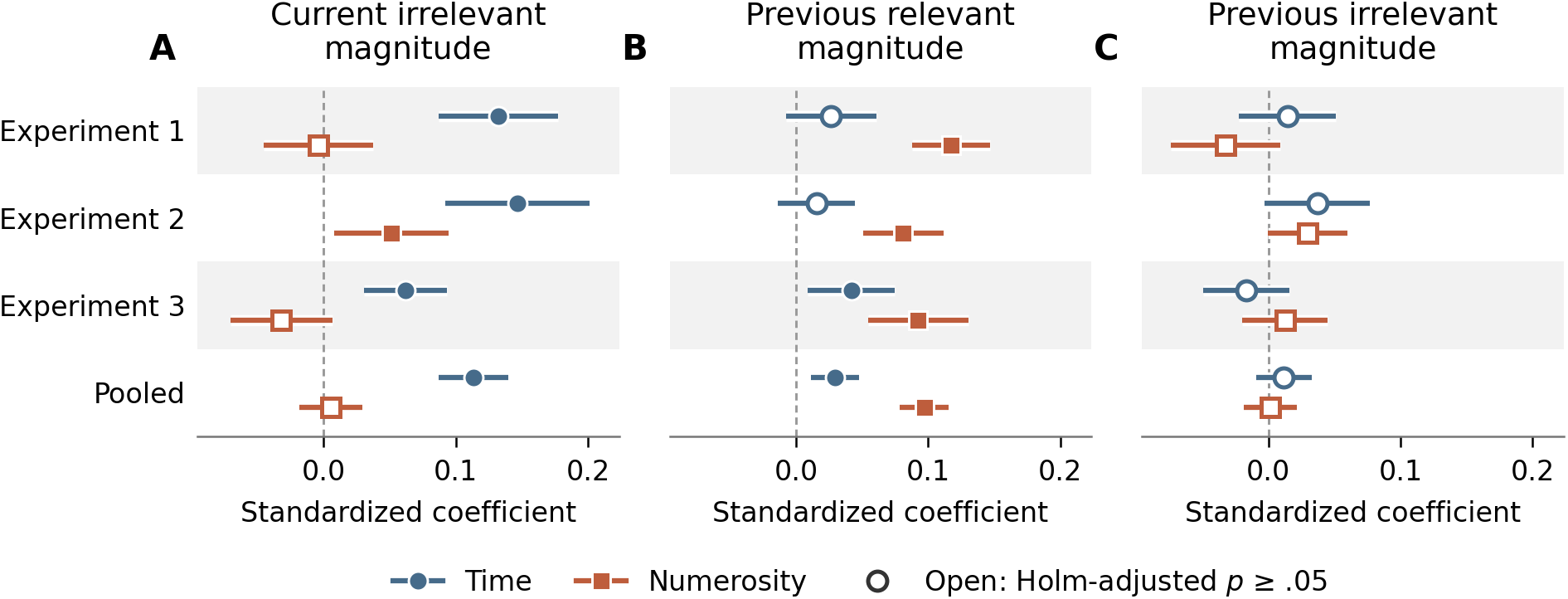
Pure blocks: within-trial integration and cross-dimensional stimulus history by experiment and current task. *Note.* (A) Current irrelevant magnitude from the integration model (current relevant and irrelevant targets). (B, C) Previous relevant and previous irrelevant magnitudes from the stimulus-history model, which adds both previous targets. Rows are the three experiments (20 participants each) and the pooled sample (60 participants). Points are standardized coefficients with participant-clustered CR2 95% CIs and Satterthwaite degrees of freedom; open markers denote Holm-adjusted *p* ≥ .05 across the two current tasks within each scope and term.

### Sensitivity of the attenuation contrasts

Time attenuation was preserved under common-scale and raw-centered sensitivities. Dots attenuation retained the same negative direction but was less stable when control paths varied by context: the common-scale result remained negative, *p* = .049, whereas the raw-centered result was uncertain, *p* = .073 (Supplementary Section S3). The combined evidence thus supported weaker carryover after task switches, with greater sensitivity to model specification for Dots.

## General Discussion

Serial dependence is usually framed as a bias from recent perceptual and decisional history. Magnitude reproduction introduces another potential source of history. Each trial contains not only a presented magnitude but also a continuously produced report. We asked whether carryover follows the magnitude that was presented, the task in which it was judged, or the response features through which it was reported. Across the three experiments, previous-response information transferred between duration and numerosity and survived task switches and changes of response key. Experiment 3 broke the fixed relation between the reported magnitude and the action used to express it by randomizing their mapping across trials, and carryover changed with that mapping. What persisted into the next judgment was report history, not only the represented magnitude.

## Against a dimension-specific account

Conventional stimulus-history attraction was clearest for numerosity repetitions, in line with the attractive serial dependence that stimulus-based measures find for numerosity (Fornaciai & Park, 2018), and it was less consistent for duration and uncertain after task switches. Once the previous response entered the model, the residual stimulus paths were small or, for duration, negative. This is the push-pull pattern described for orientation, attraction toward the previous response combined with repulsion from the previous stimulus (Fritsche et al., 2017; Gallagher & Benton, 2024; Pascucci et al., 2019; Sadil et al., 2024). It has also been found for duration, in bisection, where judgments assimilate to the previous response but shift away from the previous interval (Wiener et al., 2014), and in reproduction with the SEM approach used here (Wu, Serçe, et al., 2026). The present data extend that dissociation to numerosity reproduction and to transitions between the two magnitudes.

Our results are difficult for a purely dimension-specific account. If duration and numerosity maintained independent stimulus histories, carryover should have stayed mainly within dimension and should have run through the previous stimulus (Togoli, Fedele, et al., 2021). Cross-dimensional carryover was reliable in both directions in Experiments 1 and 2 and was captured by the previous-response path, with little evidence for a positive direct stimulus path. A response route of this kind is consistent with decisional carryover that generalizes across size and duration judgments of the same display (Wehrman et al., 2020), but the present design shows it crossing between dimensions that were judged on different trials.

This pattern also helps clarify earlier null cross-dimensional findings. In Togoli, Fedele, et al. (2021), the off-task magnitude was viewed but not reproduced. Their null result is compatible with both independent stimulus histories and the absence of a report episode for the off-task dimension. The present experiments separate these accounts. Both dimensions were actively reported.

## Where the asymmetry lives

Within a trial, numerosity biased duration reproduction in all three experiments, whereas duration biased numerosity only in Experiment 2. Earlier work has found both asymmetric and bidirectional interference (Dormal et al., 2006; Javadi & Aichelburg, 2012; Togoli, Fornaciai, et al., 2021; Xuan et al., 2007). The report, by contrast, was symmetric by design. Both magnitudes were produced through one accumulate-and-stop action, and the history that crossed between tasks ran through that action. The two parts of the data therefore separate. Integration follows the encoding stimulus and is asymmetric; report history follows the shared production format and crossed in both directions. The pure blocks also bounded cross-dimensional stimulus history near zero. This matches the null transfer that Togoli, Fedele, et al. (2021) observed under a fixed task, and it shows that a shared encoding stage, which was active within the trial, did not by itself carry the irrelevant magnitude into the next one.

## Task set modulates but does not gate response history

The task-switch results also argue against a strict task-set account. If history were retrieved only within the currently cued task, switching between duration and numerosity should have abolished the response path, yet response-history carryover remained reliable on switch trials for both current dimensions. Participants clearly processed the task switch. In Experiment 2, switch trials produced cue-to-press costs of about 40–47 ms, and wrong-key attempts were rare. Task selection changed behavior measurably, yet response-history effects persisted across switches.

Switching reduced response-history carryover. In the pooled analysis, carryover was weaker after task switches than after repetitions within mixed blocks for both current Time and current Dots judgments. The attenuation was numerically larger for Time, but the direct comparison between dimensions was uncertain. Task set modulated the strength of response history without acting as an all-or-none gate, consistent with views in which attention and task context regulate serial dependence (Kiyonaga et al., 2017). The pattern parallels sequence effects in action control, where bindings between stimulus features and responses are retrieved across a task switch even though task preparation changes other sequential effects (Lamy et al., 2024; Toledano et al., 2026). It also qualifies the finding that switching the responding hand weakens the serial dependence of confidence ratings (Bocheva & Rahnev, 2025). In Experiments 2 and 3 the key changed with the task, and carryover weakened but survived, so the feature that recurred was the shared accumulate-and-stop report rather than the key. A contrasting boundary comes from duration judgments across sensory modalities, where decisional carryover disappears when the previous and current stimuli differ in modality although the judged magnitude is the same (Li et al., 2023). What must recur for retrieval is therefore neither the task label nor the magnitude but a sufficient share of the episode’s features.

## A binding-and-retrieval account

Event-file theories propose that a trial binds stimulus, task, and response features into an episode, and that later events retrieve features of that episode when they recur (Frings et al., 2020; Hommel et al., 2001; Hommel, 2004). This framework provides a natural route for cross-dimensional carryover when response structure is preserved even as the judged dimension changes. This condition held in the present procedure. Whatever the cue, participants held a key while the display evolved and released it when the produced value matched memory. Experiment 2 constrains the identity of this episode. Time and Dots reports used different keys. Reliable cross-dimensional carryover under separate keys cannot be reduced to repetition of the same key or its motor trace. This points to a broader response episode as the relevant unit of carryover.

Experiment 3 tested the prediction that separates a shared-magnitude account from a binding-and-retrieval account by separating reported magnitude from response duration on Dots-to-Time transitions. If response history carried only a common magnitude code (Fornaciai et al., 2026), its relation to the reported value should have been preserved across production mappings. An effect tied to response production should instead vary when that mapping changes. The influence of the previous report changed with the randomized mapping, showing that response history was not determined by represented magnitude alone. This result does not rule out shared magnitude computations, which can produce transfer from a magnitude that was only viewed (Fornaciai et al., 2026) or from a task-irrelevant numeral (Fornaciai & Park, 2022). It shows that once a magnitude has been reported, the way it was produced also shapes what the next trial retrieves.

The mapping result also constrains what the episode contains. In most binding paradigms, bindings connect discrete stimulus and response categories and are inferred from partial-repetition costs. The relevant production information here was continuous. The previous numerosity report was always made with the same key, so it had no distinct response identity of the kind that binding paradigms track. What differed between trials was how long that key was held to report a given number, because the randomized mapping made a large number require a long hold on some trials and a short hold on others. Carryover depended on the mapping between reported number and hold duration. The history retained from a report therefore includes how the report was produced, not only the value it delivered. This extends the relevance of event-file logic beyond discrete response identities to continuous features of response production, in line with evidence that serial dependence scales with the depth of action binding in duration reports (Wu, Gündogdu, et al., 2026).

The context analyses qualify this picture. Response history was not carried forward with fixed strength. Carryover was weaker after a task switch than after a repetition within the same mixed-block context, and Time carryover differed even between repetitions in mixed and pure blocks. What a previous response episode did to the current judgment depended on what had been reported, on how it had been produced, and on the context in which that history was encountered again. In binding-and-retrieval terms, retrieval appears sensitive to the match between the current event and the stored episode, as in visual working memory, where serial dependence is stronger between objects that share context features across trials (C. Fischer et al., 2020).

Binding and retrieval describe how information from a previous event becomes available again. They say less about how strongly that information shapes the current judgment. Previous representations can be reactivated by current input (Bae & Luck, 2019), and reactivation of prior reports has been linked to subsequent serial bias (Zhang & Lewis-Peacock, 2024). The weight of recent history need not be fixed. In temporal reproduction it varies with current evidence quality and, more tentatively, contextual stability (Qu & Shi, 2026), and individual differences in such biases track beliefs about whether successive stimuli are temporally continuous (Glasauer & Shi, 2022). Serial dependence has been located after perception, in working memory (Bliss et al., 2017; Czoschke et al., 2019). Our findings place one carrier a stage later still, at the production of the report. Several studies have shown that reporting can strengthen attractive serial dependence and that carryover depends on the response procedure: numerosity serial dependence is stronger after a reported than a merely viewed stimulus (Morimoto & Makioka, 2024), duration carryover follows active reproduction but not passive viewing (Cheng, Chen, Glasauer, et al., 2024), the previous interval biases reproduction only when it was tied to a motor command (Wehrman et al., 2018), and decisional carryover in duration vanishes when the response mode changes (Wu, Serçe, et al., 2026). The mapping result adds that its production features are part of what is retrieved. The same previous number pulled the next duration judgment toward longer values when it had been reported with a long hold and not when it had been reported with a short hold, and this influence crossed from number to time and survived a change of task. The conventional analysis, which indexes history by the previous stimulus, found no cross-task serial dependence in any experiment; the carrier became visible only once the previous response entered the model. An attractive lag-1 effect therefore need not be the persistence of a stimulus or a decision. It can be the trace of the action through which the previous judgment was expressed, and studies that measure serial dependence from the stimulus alone will miss it. Identifying serial dependence requires asking which part of the past is carried forward, not only how strongly.

## Limitations

The mapping manipulation in Experiment 3 was directional. Ascending and descending reports differed not only in the hold needed to reach a given number but also in the direction of the visible change, so the two mappings reversed the value-to-hold relation and the visual sequence together. Random assignment licenses attributing the change in carryover to the mapping, but not to one of its consequences, whether motor, temporal, visual, or decisional. A non-monotonic modulation of the visible count, for example a sinusoidal one in which the same number is reached at several hold durations, would separate hold duration from display direction. The test was also one-sided. It was decisive for Dots-to-Time transitions, whereas on Time trials reported duration and hold duration coincide, so the reverse transition afforded no comparable test.

The encoding display was static. This choice kept the within-trial time–number asymmetry in place where it could be measured, but it also limits the integration results to arrays whose number does not change while they are viewed. More symmetric interference has been reported for judgments of the duration and average numerosity of dot-array sequences (Togoli, Fornaciai, et al., 2021), and whether response history behaves the same way after such encoding remains to be tested.

The remaining limits concern generalization. Each experiment retained 20 participants from a local university population, with demographics limited to age and self-reported gender. The accumulate-and-stop procedure gave both dimensions the same additive response structure, which may itself be part of why history transferred between them. Whether similar transfer appears with bidirectional adjustment, verbal report, or discrete choice remains open.

## Conclusion

Three experiments asked what a preceding report leaves behind. Its influence crossed changes in magnitude dimension, task, and physical key, weakened after a task switch, and changed with the randomized relation between a reported number and the hold that produced it. A binding-and-retrieval account explains this pattern: each report binds the reproduced magnitude to the action, task, and key that produced it, and the next trial retrieves that episode to the extent that its features recur. The experiments do not identify which feature of production carried the effect. What they show is that serial dependence carries the history of a previous report episode, not only of the preceding stimulus or the value it represented.

