## Supplemental Files for "Serial dependence carries the action that produced the previous report"

### Overview

The supplementary sections follow the order in which the main text cites them. Section S1 reports the conventional and response-adjusted stimulus-history regressions. Section S2 reports stimulus-report fidelity and the previous-stimulus control path. Section S3 reports the task-switch contrasts and their sensitivity to scaling and control paths. Section S4 reports the Experiment 2 task-key checks, and Section S5 the Experiment 3 randomized-mapping models.

Machine-readable tables and reproducible code accompany the submission repository.

### S1. Conventional and response-adjusted stimulus history

#### Question and model

The conventional analysis estimated previous-stimulus serial dependence before conditioning on the previous response. Separate regressions were fitted for each experiment, current task, and transition context, with current bias regressed on the previous stimulus, the current stimulus, and participant fixed effects. Matched response-adjusted regressions added the previous response feature. All variables used the exact complete cases, participant × context centering, and within-context standardization of the primary context SEM. On switches, both stimulus predictors were in the current judgment dimension. Current- and previous-stimulus coefficients were free to vary across contexts in these additional regressions; the primary coordinated SEM retained its original common-control specification.

On switch trials, the previous stimulus and response concerned different dimensions, so we did not combine the paths as $c' + ab$. These analyses describe associations and do not establish causal mediation.

**Result**

Figure S1 shows the conventional coefficient $c$ and the response-adjusted coefficient $c'$ for every experiment, current task, and transition context. Conventional attraction was clearest for Dots repetitions. Pure-block Dots estimates survived Holm correction in all three experiments ($c$ = 0.117, 0.081, and 0.094), as did the mixed-repetition Dots estimates in Experiments 2 and 3 (0.105 and 0.073). Pure-block Time attraction survived correction only in Experiment 3 ($c$ = 0.045); the Time estimates in Experiments 1 and 2 and all stimulus-only switch estimates were uncertain after correction.

After adjustment for the previous response, the Time coefficient was negative in pure blocks in all three experiments ($c'$ = −0.098, −0.086, and −0.057) and on mixed repetitions in Experiment 3 (−0.080). For Dots, pure-block stimulus history remained positive in Experiment 3 ($c'$ = 0.052). The other response-adjusted tests did not survive correction. Conditioning on the previous response therefore did not produce a uniform negative stimulus coefficient across tasks and experiments.

**Figure S1**

*Conventional and response-adjusted stimulus history by experiment and transition context*

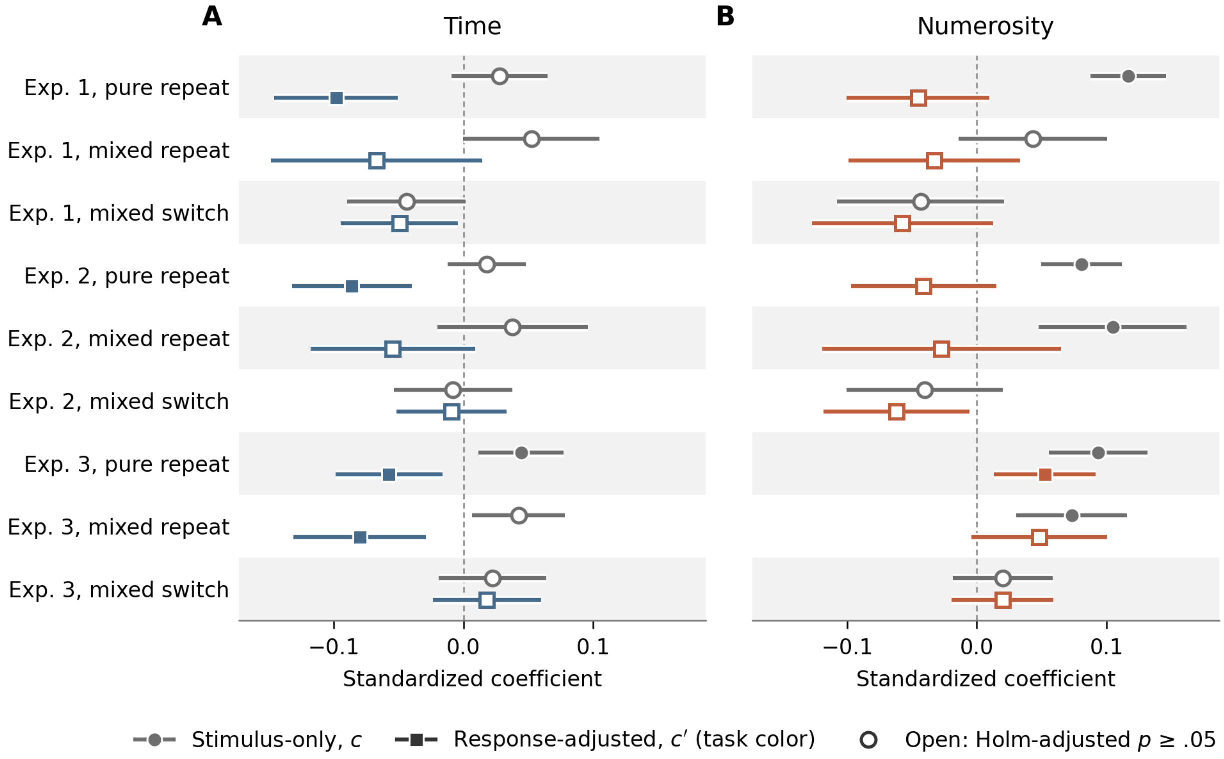


*Note.* Panel A shows Time judgments and Panel B Numerosity judgments. Rows are the three transition contexts within each experiment. Gray circles show the stimulus-only coefficient $c$; colored squares show the response-adjusted coefficient $c'$ from the matched regression on the same trials. Error bars are participant-clustered CR2 95% CIs with Satterthwaite degrees of freedom; open markers denote estimates whose Holm-adjusted $p$, over the six task-by-context tests per experiment and coefficient, was at or above .05. Every cell included 20 participants. Variables were centered within participant × context and standardized within context. These regressions allowed both stimulus slopes to vary across contexts; the primary response-history SEM constrained those paths.

### S2. Stimulus-report fidelity and stimulus-control paths

Path *a* relates the previous target to its report and, like *b*, takes a separate value in each transition context; because it concerns the previous task, on switches it describes the report of the other dimension. Path *c*′ relates the previous stimulus in the current judgment dimension to current bias, conditional on the previous response feature and the current stimulus, and was held equal across contexts; the matched regressions in Section S1 relax this constraint. Fidelity was high in every context (Figure S2). Path *a* ranged from 0.459 to 0.688 across experiments, tasks, and contexts, and the switch values (0.585–0.648 for current Time, 0.466–0.517 for current Dots) matched the fidelity of the other task's report. The constrained previous-stimulus path was negative for Time in all three experiments, *c*′ = −0.074, 95% CI [−0.105, −0.044]; −0.051, [−0.080, −0.022]; and −0.040, [−0.065, −0.014], and for Dots in Experiments 1 and 2, −0.049, [−0.086, −0.013] and −0.046, [−0.080, −0.012], whereas it was positive for Dots in Experiment 3, 0.041, [0.017, 0.064]. In the combined sample, the path was −0.054, [−0.071, −0.037] for current Time and −0.012, [−0.032, 0.007] for current Dots.

**Figure S2**

*Stimulus-report fidelity by experiment and transition context*

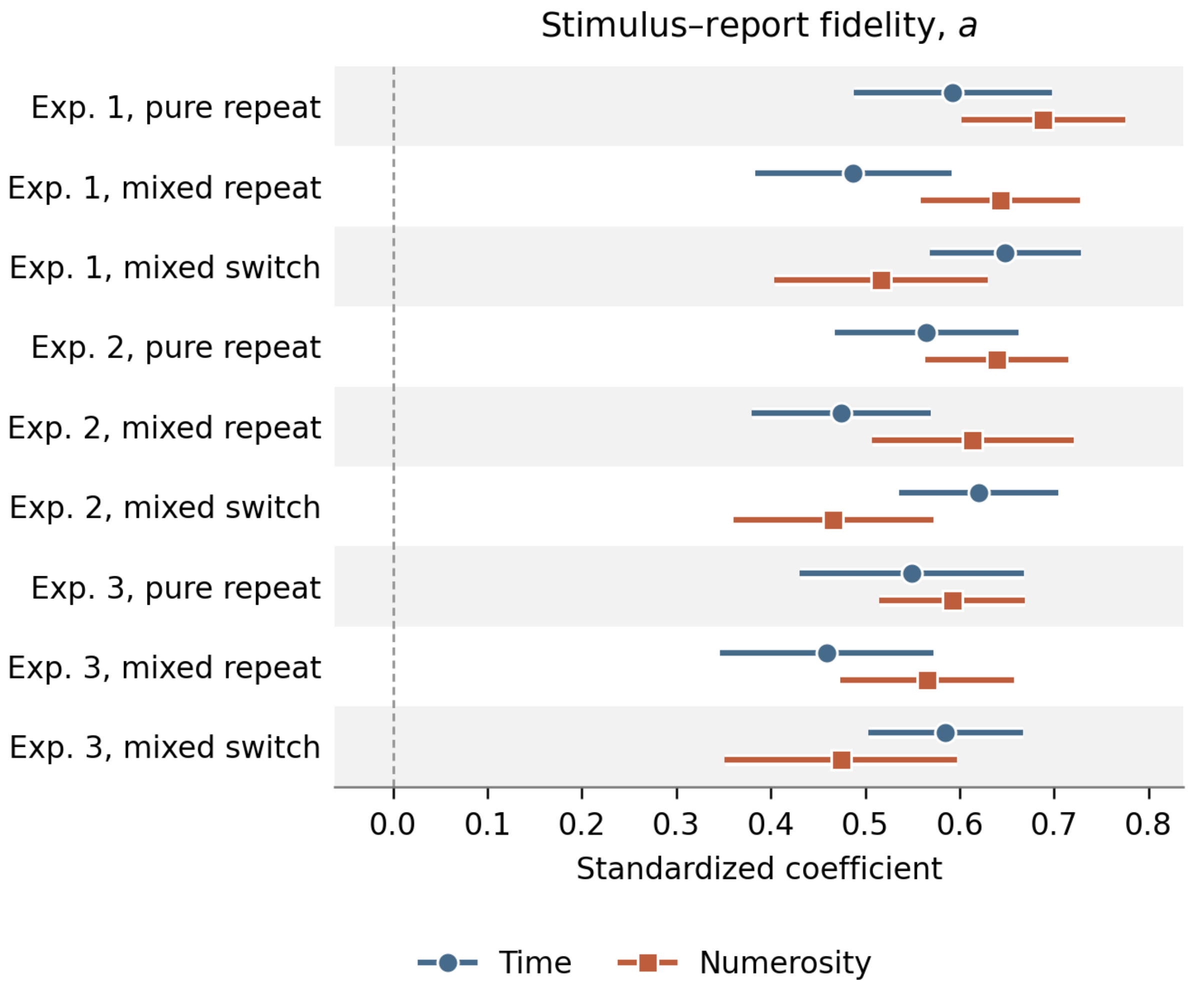


*Note.* Rows are the three transition contexts within each experiment; Time and Numerosity judgments are dodged within rows. Points are the fidelity path *a* from the response-history model, with 95% CIs from participant-clustered MLR inference. On switches the path concerns the other task's report: the previous numerosity report for current Time and the previous duration report for current Dots. Variables were centered within participant × context and standardized within context.

### S3. Task switching and response-history carryover

#### Question and model

Did switching reduce carryover relative to repeating a task within mixed blocks? Planned differences used a stacked participant-fixed-effect regression with participant-clustered CR2 covariance. It reproduced the coordinated model's coefficients and contrasts to within 0.00136. Within each experiment and in the combined sample, the six task-by-contrast tests (two current tasks × three context differences) formed one Holm family; the direct Time-minus-Dots comparison was tested separately.

#### Result

Figure S3 shows the three context contrasts for each experiment and for the combined sample. Across experiments, switching attenuated carryover for both current tasks (combined Time $\Delta b$ = −0.110, 95% CI [−0.158, −0.062]; Dots $\Delta b$ = −0.053, [−0.094, −0.011]). Attenuation was numerically larger for Time, but the direct Time-minus-Dots difference was uncertain, $\Delta b$ = −0.057, 95% CI [−0.119, 0.005], $p$ = .069; this contrast used 33,415 trials from 60 participants. The Holm adjustment retained three Experiment 1 contrasts, the Time contrast in Experiment 3, and all five nominally reliable combined-sample contrasts; neither nominal Experiment 2 contrast survived. In the combined sample, mixed switches also showed weaker carryover than pure repetitions for Time, $\Delta b$ = −0.058, 95% CI [−0.095, −0.020], $p$ = .003, and for Dots, $\Delta b$ = −0.090, [−0.128, −0.051], $p$ < .001. Because these contrasts combine a task change with a change in block context, they do not isolate either factor.

**Figure S3**

*Transition-context contrasts of response-feature carryover*

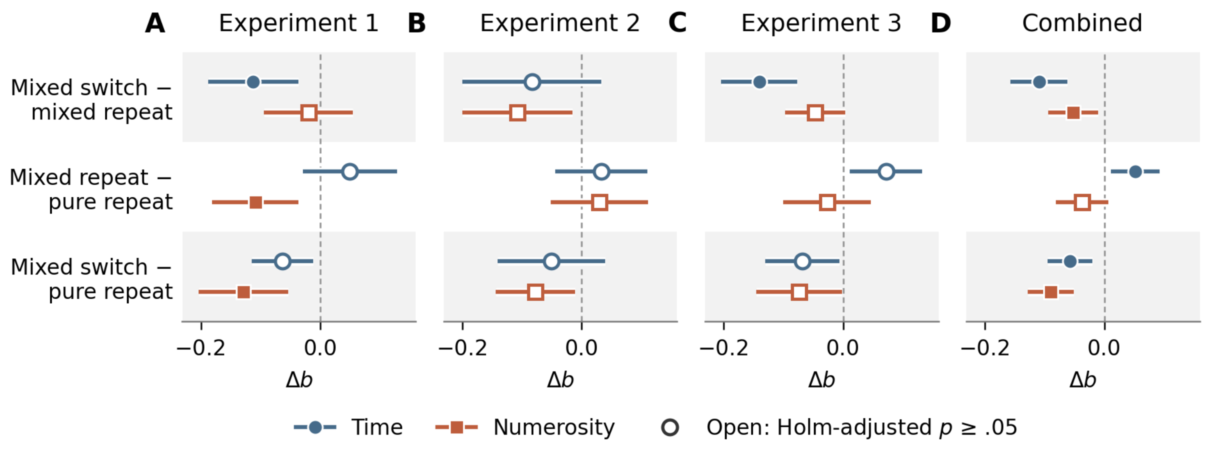


*Note.* Panels A–C show Experiments 1–3 and Panel D the combined sample. Rows are the three planned differences in the carryover path *b*; negative values indicate weaker carryover in the first-named context. Points are standardized differences with participant-clustered CR2 95% CIs and Satterthwaite degrees of freedom; open markers denote Holm-adjusted $p \geq .05$ within the six-test family of each panel. Each experiment-specific cell contained all 20 retained participants.

### Trial-level trends

Figure S4 shows the trial-level trends underlying the context-specific carryover coefficients in main-text Figures 3–5 (Panel A), together with the combined-sample trends.

**Figure S4**

*Response-history trends across pure repetitions, mixed repetitions, and mixed switches*

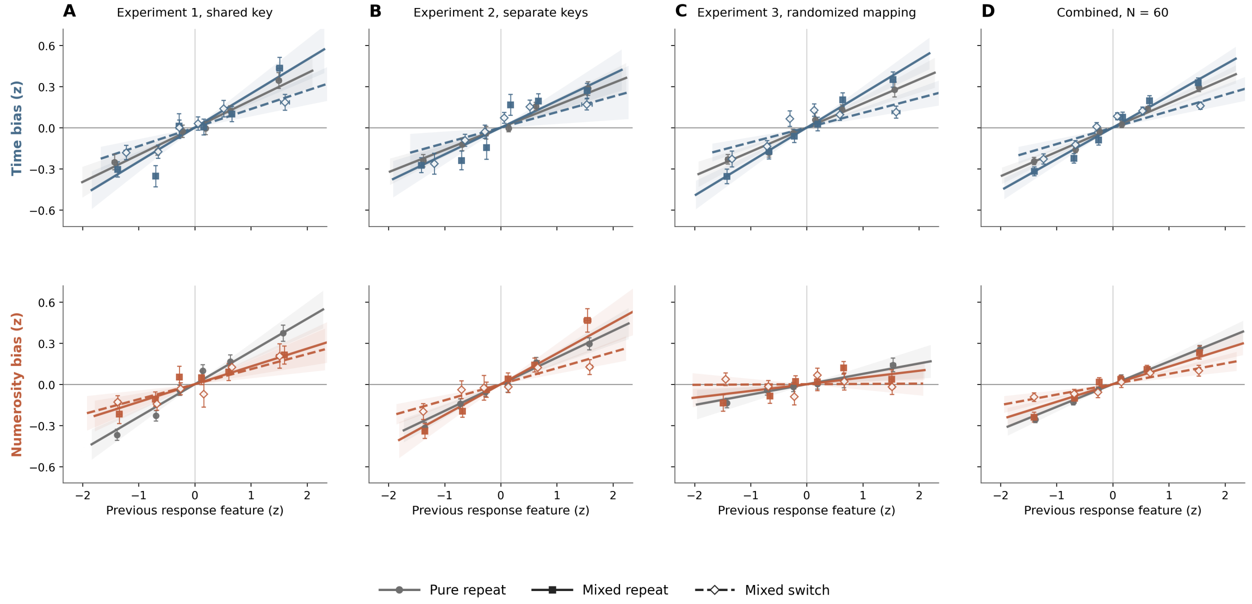


*Note.* Panels A, B, and C show Experiments 1, 2, and 3, and Panel D shows the combined sample; within each panel, Time is shown above Numerosity. Points are participant-first sextile means and SEMs after previous- and current-stimulus contributions were removed from the outcome. Lines show the context-specific response-history path; bands are 95% CIs. On switches, the previous response feature was hold duration for current Time judgments and endpoint count for current Numerosity judgments.

### Scaling and control-path sensitivity

Time attenuation was stable across four alternative specifications, which crossed common-scale or raw-centered variables with stimulus-control paths that were either constrained across contexts or free to vary ($\Delta b$ between −0.131 and −0.149, all $p < .001$). The Dots estimate had the same direction throughout ($\Delta b$ between −0.038 and −0.059) and remained reliable when the control paths were constrained ($p = .007$ with the common scale and .012 raw centered); it was less precise when those paths varied by context ($p$ = .049 with the common scale and .073 raw centered). Common-scale models used one experiment-by-task standard deviation across contexts; raw-centered models applied no standard-deviation division. All models retained participant-by-context centering. Context-standardized coefficients are standardized associations, not scale-invariant percentage changes.

### S4. Experiment 2 task-key compliance

Wrong-key attempts occurred on 69 of 11,520 scheduled main trials (0.60%), with similar rates on Time (0.66%) and Dots (0.54%) trials. In mixed blocks, rates were 0.43% on repetitions and 1.80% on switches, and were similar across key counterbalances (0.57% and 0.63%). Correct-key cue-to-press latencies showed switch costs for Time, 40.47 ms, 95% CI [12.58, 68.36], $t(19) = 3.04$, $p = .007$, $d_z = 0.68$, and Dots, 47.25 ms, 95% CI [27.52, 66.98], $t(19) = 5.01$, $p < .001$, $d_z = 1.12$. Participants used the task keys as instructed and incurred a measurable switching cost.

### S5. Randomized response mapping in Experiment 3

Models of current reproduction bias included a separate intercept for each participant and each combination of previous and current mapping. This adjustment prevented stable differences in the effect of starting count from entering the mapping-specific slopes.

#### Coordinate sensitivity

The main-text model estimated an uncertain mapping-averaged numerosity coefficient, $b = 0.054$, 95% CI [−0.020, 0.128], $p = .146$, and a positive numerosity-by-mapping interaction, $b = 0.090$, 95% CI [0.051, 0.128], $p < .001$. With a fixed-design-mean ratio, the mapping-averaged coefficient was 0.046, 95% CI [−0.018, 0.111], $p = .146$, and the interaction

0.077, [0.044, 0.111], $p < .001$; with a log ratio, the two were 0.063, [−0.012, 0.138], $p = .092$, and 0.067, [0.028, 0.105], $p = .002$. Fixed-design-mean estimates need not equal the standardized main-text coefficients. Re-centering the ratio predictors within participant did not change the focal slopes or their tests, so we do not report a separate participant-centered analysis.

**Joint observed-trace model**

The joint model entered previous reported numerosity and hold duration simultaneously. Hold duration retained a positive conditional association with current Time bias in both coordinates, whereas numerosity did not. With the fixed-design-mean ratio, hold duration given numerosity was $b = 0.095$, 95% CI [0.060, 0.131], $p < .001$; numerosity given hold duration was 0.043, [−0.020, 0.106], $p = .170$; and their covariance-aware difference was 0.052, [−0.023, 0.127], $p = .158$. With the log ratio, the corresponding values were 0.086, [0.037, 0.134], $p = .002$; 0.055, [−0.014, 0.124], $p = .110$; and 0.030, [−0.038, 0.099], $p = .361$. Both trace coefficients varied with current mapping and were constrained across previous mapping. The analysis does not establish that hold duration has the larger coefficient.

Adding the hold-duration block improved fit, $F(2, 15.68) = 17.33$, $p < .001$. A direction-saturated model, in which both coefficients varied with previous and current mapping, could not separate the two readouts. Its hold coefficient was positive but imprecise, $b_h = 0.084$, 95% CI [−0.042, 0.210], $p = .178$, and its difference from numerosity was uncertain, $b_h - b_r = 0.013$, 95% CI [−0.154, 0.180], $p = .872$.

To compare trials with the same reported numerosity, the model treated each reported number as a category. The stricter version allowed these number effects to differ by previous mapping. Time bias increased with hold duration in the first model, $b = 0.098$, 95% CI

[0.047, 0.149], $p < .001$. The stricter version was less precise, $b = 0.102$, 95% CI [−0.020, 0.225], $p = .095$.

Next, we separated the part of hold duration induced by the randomized mapping from the variation remaining within each mapping. The first stage predicted standardized hold duration from reported numerosity, previous mapping, their interaction, and the previous stimulus. A four-degree-of-freedom spline allowed the effect of reported numerosity to be nonlinear. The model also included a separate intercept for each participant and previous mapping. For each trial, the mapping-induced component was half the difference between predicted hold duration under the observed and opposite mappings. The within-mapping component was the first-stage residual. Both components were standardized.

The second stage predicted standardized current Time bias from both components, reported numerosity, and the previous and current stimuli. It included a separate intercept for each participant and each combination of previous and current mapping. The mapping terms accounted for 80.1% of the remaining hold-duration variance (partial $R^2 = .801$). The mapping-induced component predicted current Time bias, $b = 0.084$, 95% CI [0.033, 0.135], $p = .003$. The within-mapping component was uncertain, $b = 0.034$, 95% CI [−0.017, 0.085], $p = .175$. A participant-cluster bootstrap repeated both stages in each of 2,000 resamples. It gave $b = 0.085$ with a percentile 95% CI [0.040, 0.138].

**Reverse transition**

For Time → Dots, the previous endpoint had a near-zero mapping-averaged slope, $b = -0.010$, 95% CI [−0.062, 0.043], $p = .702$. Its interaction with previous and current mapping was also uncertain, $b = 0.005$, 95% CI [−0.050, 0.059], $p = .850$, and none of the four conditional slopes was reliable: −0.050, 95% CI [−0.171, 0.072] for descending → descending; −0.053, [−0.148, 0.042] for descending → ascending; 0.024, [−0.065, 0.112] for ascending → descending; and 0.040, [−0.063, 0.144] for ascending → ascending (all $p \geq .255$). These estimates use participant × mapping-cell intercepts, participant-clustered CR2 covariance, and Satterthwaite degrees of freedom.

Averaged over mappings, current Dots bias increased with the previous reproduced duration, $b = 0.065$, 95% CI [0.014, 0.115], $p = .015$. On Time trials, reproduced duration equals key-hold duration, so their contributions to carryover cannot be separated.
